# COP I and II dependent trafficking controls ER-associated degradation in mammalian cells

**DOI:** 10.1101/2022.04.07.487480

**Authors:** Navit Ogen-Shtern, Chieh Chang, Haddas Saad, Niv Mazkereth, Chaitanya Patel, Marina Shenkman, Gerardo Z. Lederkremer

## Abstract

Misfolded proteins and components of the endoplasmic reticulum (ER) quality control and ER associated degradation (ERAD) machineries concentrate in mammalian cells in the pericentriolar ER-derived quality control compartment (ERQC), suggesting it as a staging ground for ERAD. By tracking the chaperone calreticulin and an established ERAD substrate, asialoglycoprotein receptor H2a, we have now determined that the trafficking to the ERQC is reversible and recycling back to the ER is slower than the movement in the ER periphery. The dynamics suggest vesicular trafficking rather than diffusion. Indeed, using dominant negative mutants of ARF1 and Sar1 or the drugs Brefeldin A and H89, we observed that COPI inhibition causes accumulation in the ERQC and increased retrotranslocation and ERAD, whereas COPII inhibition has the opposite effect. Our results suggest that targeting of misfolded proteins to ERAD involves COPII-dependent transport to the ERQC and that they can be retrieved to the peripheral ER in a COPI-dependent manner.

## Introduction

During secretory protein biogenesis, after translocation into the ER, the quality control machinery facilitates protein folding and promotes segregation of misfolded proteins destined for ERAD. We had previously determined that misfolded proteins destined to ERAD accumulate in a specialized pericentriolar compartment, the ER-derived quality control compartment (ERQC), a staging ground for retrotranslocation and targeting to ERAD ^1–5^. The ERAD-associated luminal lectin OS-9 always appears localized mainly at the ERQC, whereas other quality control and ERAD components are recruited upon misfolded protein accumulation (ER stress). This compartmentalization depends on the PERK pathway of the unfolded protein response (UPR) and on the upregulation of Herp ^4^. PERK is activated by the initial accumulation of misfolded proteins and consequently upregulates Herp, resulting in a dynamic protein recruitment to the ERQC. Herp recruits the membrane-bound ubiquitin ligase HRD1 and additional quality control and ERAD machinery components, such as the luminal calreticulin (CRT), and the membrane proteins calnexin (CNX), Derlin-1, EDEM1, ERManI and cytosolic components, including the ubiquitin ligase SCFFbs2 and the AAA ATPase p97 ^2,6,7^. However, other machinery proteins such as BiP, PDI, UGGT and ERp57, do not accumulate in the ERQC^2–5,8,9^. Here we show that compartmentalization of ERAD substrates in the ERQC, and subsequent targeting to ERAD, depend on a mechanism that is consistent with vesicular trafficking, involving coat protein complex II (COPII), and that recycling from the ERQC requires COPI. COPI and COPII coats had first been identified as complexes that take part in vesicular trafficking between the ER and the Golgi ^10–12^. Yet, COPII was recently found to participate in other processes taking place from the ER as well, including the formation of ER whorls in mammalian cells ^13^ and an autophagy process targeting the ER (ER-phagy) in yeast ^14,15^.

## Results

### Dynamics of concentration of ER quality control and ERAD components at the ERQC upon proteasomal inhibition

We had previously shown that upon proteasomal inhibition with diverse inhibitors, or under other conditions in which ERAD substrates accumulate (ER stress), several ER quality control and ERAD factors concentrate at the ERQC. This can be seen, for example, by comparing the localization of an established ERAD substrate, the uncleaved precursor of asialoglycoprotein receptor H2a, linked to monomeric red fluorescent protein (H2a-RFP), which serves as a marker of the ERQC ^3,4^, with that of the soluble luminal chaperone BiP in NIH 3T3 cells. Whereas H2a-RFP was recruited to the ERQC under proteasomal inhibition with lactacystin (Lac), BiP was not (Fig. 1A), as we had previously seen ^16^. H2a-RFP accumulates in the centrosomal region of the cell but with no colocalization with a Golgi marker, GalT-YFP ^2,3^ (Fig. 1A). In contrast to BiP or BiP linked to GFP (BiP-GFP) (Fig. 1, Fig. suppl. 1), another soluble luminal chaperone, CRT linked to GFP (CRT-GFP) was recruited to the ERQC under proteasomal inhibition (Fig. 1B). A proteasomal subunit linked to YFP, PSMD14-YFP, was also partially recruited to the ERQC region upon proteasomal inhibition (Fig. 1C), similarly to what we had seen with other cytosolic ERAD factors, p97 and SCFFbs2 ^3,7^. PSMD14-YFP even showed a higher presence in the ERQC region than H2a-RFP and CRT-GFP in untreated cells (Fig. 1D). Using another approach to analyze the compartmentalization, iodixanol gradients optimized to separate the different ER-like densities ^4,6^, we observed H2a-RFP peaking in middle fractions (5-6), with a small shift to heavier fractions, peaking in fraction 7, upon proteasomal inhibition, in this case with a combination of MG132 and ALLN (Fig. 1E). We had previously characterized this shift as occurring upon protein concentration in the ERQC ^4,6^. CRT-GFP had a similar behavior and also endogenous CNX. In contrast, endogenous BiP appeared in the light fractions and its migration did not change upon proteasomal inhibition. Consistent with the fluorescence of PSMD14-YFP, most of the endogenous proteasomes, likely membrane-bound, detected with antibodies against the 19S and 20S proteasomal subunits, also appeared in the middle fractions, with a lesser shift upon proteasomal inhibition, in this case with the proteasome inhibitor bortezomib (Bz) (Fig. 1F). Therefore, using different approaches and various proteasome inhibitors we can observe accumulation in the ERQC of an ERAD substrate, proteasomes and some chaperones at the ERQC.

**Figure 1.**
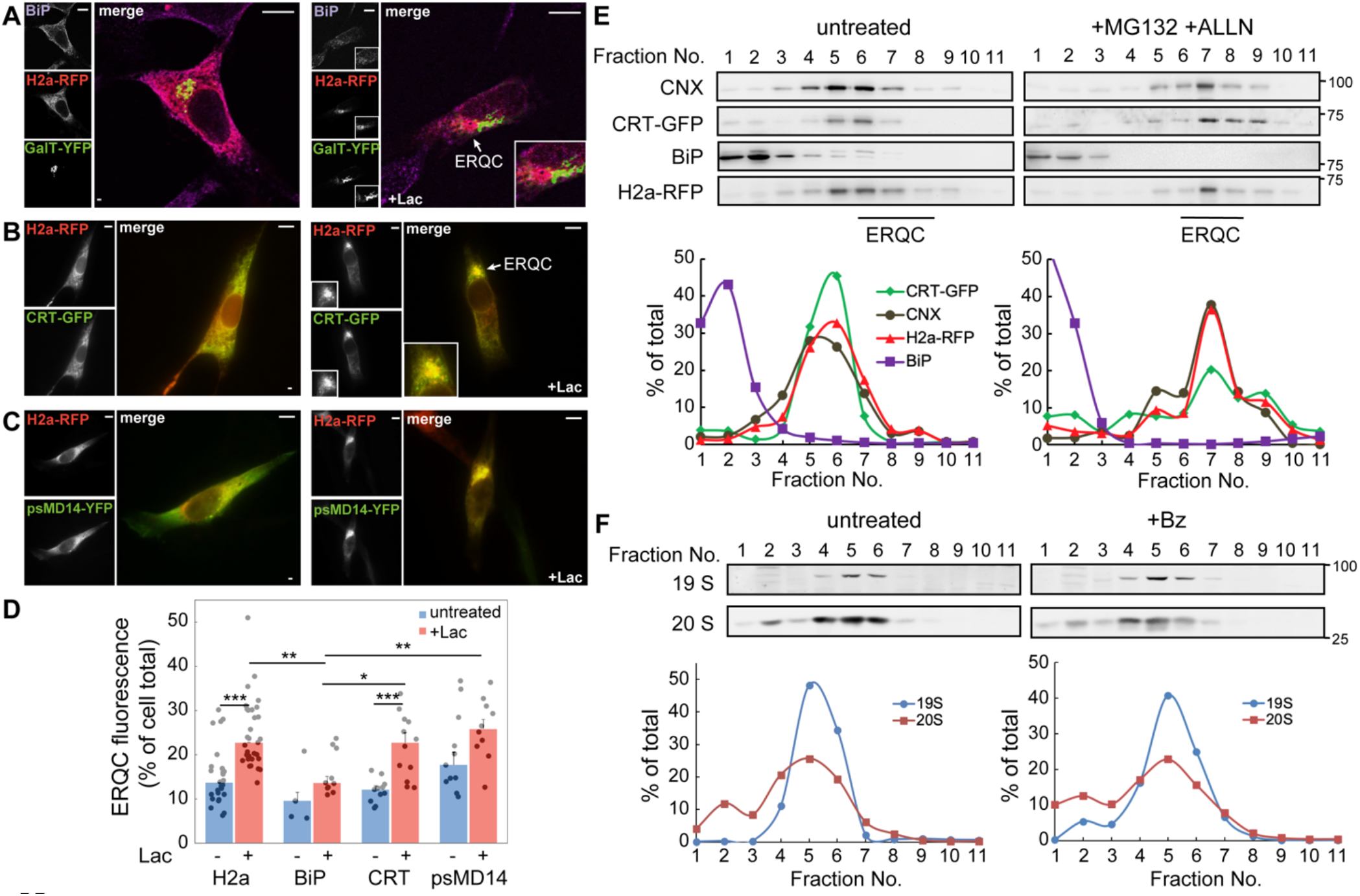
ERAD substrate H2a-RFP, CRT-GFP and proteasomes accumulate at the ERQC upon proteasomal inhibition. **(A-C)** Fluorescence microscopy of NIH 3T3 cells treated without or with the proteasomal inhibitor Lactacystin (Lac, 20μM, 3h), transiently expressing H2a-RFP and the Golgi marker GalT-YFP (A), CRT-GFP (B), or psMD14-YFP (C). CRT-GFP (B) and the proteasomal subunit psMD14-YFP (C) concentrate and colocalize with H2a-RFP at the juxtanuclear ERQC, especially in Lac-treated cells. In contrast, there is no colocalization of H2a-RFP with GalT-YFP and no concentration of BiP in Lac-treated cells (A). Bars= 10μm. **(D)** The percentage of fluorescence in the ERQC compared to the entire cell was measured in the cells in (A-C). The graph shows mean +-SEM of cells from three independent experiments (10 to 30 cells per population). P value +Lac vs. untr.: H2a-RFP= 4.8 x 10^-6^, CRT-GFP= 0.00027; vs. BiP+Lac: H2a-RFP= 0.0015, CRT-GFP= 0.029, psMD14-YFP= 0.0013. **(E)** Immunoblots of fractions of iodixanol density gradients from NIH 3T3 cells transiently expressing CRT-GFP and H2a-RFP show comigration of CRT-GFP, H2a-RFP and endogenous CNX in mid-density fractions (5-7), shifting to denser fractions (6-8) (ERQC) upon MG-132+ALLN (50μM, 3h) treatment, compared to untreated cells. BiP appears in lighter fractions and does not shift upon proteasomal inhibition. MW markers in kDa are indicated on the right. The graphs show quantification of the percentage of the signal in each fraction relative to the sum of all fractions. The experiment is representative of 3 independent repeat experiments. **(F)** In a similar experiment, most of the endogenous proteasome subunits 19S and 20S appear in similar mid-density fractions upon iodixanol fractionation. In this case proteasomal inhibition was done for 3h with Bz (1μM). Representative of 3 independent repeat experiments.

To analyze the dynamics of protein concentration at the ERQC, we performed a live cell time-lapse experiment with cells expressing H2a-RFP and CRT-GFP. In this case, we used CHO cells, which are less mobile, to facilitate measurements for several hours. A stable cell line expressing CRT-GFP was established, which was transfected with H2a-RFP and treated with Lac for 3 h, during which we conducted live cell imaging (Fig. 2A). Both H2a-RFP and CRT-GFP concentrated in the ERQC during this period, although a larger portion of CRT-GFP remained in the peripheral ER compared to H2a-RFP (Fig. 2A, B). H2a-RFP concentrated in the ERQC at a rate of 13% of the total cellular protein per minute, whereas the rate for CRT-GFP was 4.1% per minute, a 3.25-fold slower rate (Fig. 2B).

**Figure 2.**
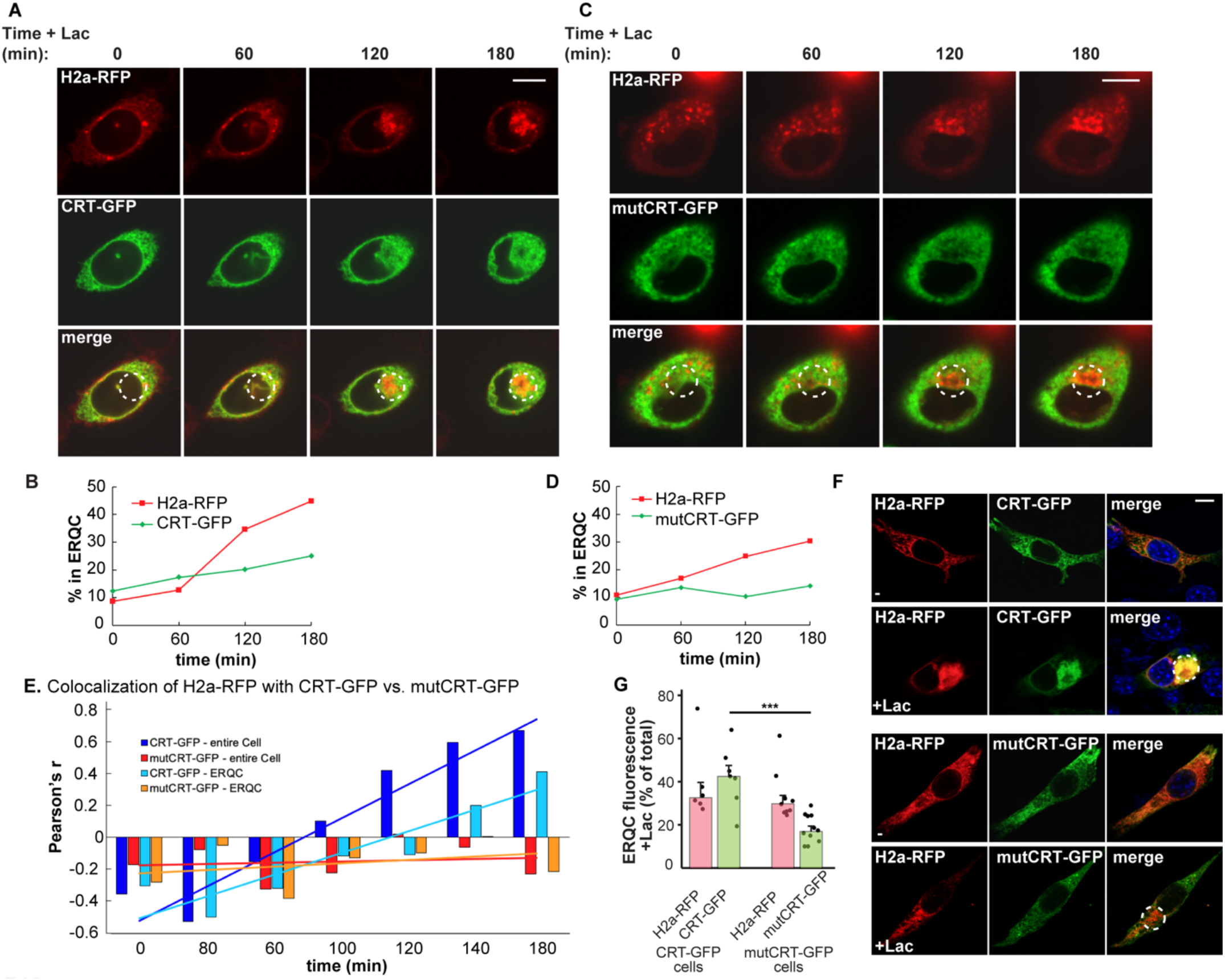
Movement of H2a-RFP and CRT-GFP to the ERQC. CRT-GFP dependence on its lectin activity. **(A)** CHO cells stably expressing CRT-GFP were transiently transfected with H2a-RFP and treated with Lac (10μM). Live cell imaging shows the dynamic concentration of both proteins in the juxtanuclear ERQC. Bar= 10μm. **(B)** The percentage of H2a-RFP and CRT-GFP concentrated in the ERQC (dashed circles in (A)) was calculated at the different time points. **(C-D)** CHO cells stably expressing mutCRT-GFP, lacking a functional lectin domain, were transfected with H2a-RFP and treated with 10μM Lac. Live cell imaging shows absence of mutCRT-GFP accumulation at the ERQC. Bar= 10μm. **(E)** Pearson’s coefficients of colocalization of CRT-GFP or mutCRT-GFP with H2a-RFP in the entire cell or in the area of the ERQC were measured at different time points in the experiments in (A) and (B). mutCRT-GFP showed no colocalization with H2a-RFP. The results are representative of 3 independent experiments. **(F)** In Lac-treated NIH 3T3 cells (20μM, 3h), CRT-GFP and H2a-RFP accumulate in the ERQC, whereas untreated cells show an ER pattern. No colocalization appears in the ERQC (dashed circles) of NIH 3T3 cells expressing mutCRT-GFP and H2a-RFP after incubation with Lac, compared to untreated cells. **(G)** The percentage of GFP or RFP fluorescence in the ERQC compared to the entire cell was measured in Lac-treated cells expressing CRT-GFP plus H2a-RFP or mutCRT-GFP plus H2a-RFP. The graph shows mean +-SEM of cells from two independent experiments (~20 cells per population). P value CRT-GFP vs. mutCRT-GFP = 0.00014.

The interaction of CRT with its glycoprotein substrates is dependent on a functional lectin domain. We established a stable cell line expressing CRT(Y108F)-GFP, which carries a point mutation in its lectin domain ^17^. A time lapse experiment similar to the one described above using WT CRT-GFP was performed with this mutant (mutCRT-GFP). While H2a-RFP concentrated in the ERQC of these cells as before, although at a slower rate (6.6% per min), mutCRT-GFP did not concentrate to any significant extent within the 3h time frame (1.1% per min) (Fig. 2C, D). Although the rate of protein concentration varied to some extent in repeat experiments, mutCRT-GFP did not accumulate in the ERQC (Fig. 2, Fig. suppl. 1).

We compared the colocalization by calculating the Pearson’s coefficient. For WT CRT-GFP the colocalization with H2a-RFP increased with time, whether only the ERQC region or the entire cell were measured (light or dark blue trendlines respectively in Fig. 2E). The colocalization measured for the entire cell area increased at a higher rate and to a higher extent than that measured for the ERQC, suggesting that colocalization of CRT-GFP and H2a-RFP already occurs in peripheral ER sites, before transport to the ERQC. In contrast, there was no colocalization and no increase with time for mutCRT-GFP (Fig. 2E, orange and red trendlines) suggesting that the binding to substrates through its lectin domain is necessary for the accumulation of CRT in the ERQC. These results were not unique to the CHO cell lines, they were similar in NIH 3T3 cells, where after 3h of Lac treatment mutCRT-GFP concentrated in the ERQC to a much lower extent than WT CRT-GFP (Fig. 2F, G).

### Recycling from the ERQC to the peripheral ER

To study the reversibility of the process of protein concentration in the ERQC, we treated cells expressing H2a-RFP and CRT-GFP with the reversible proteasome inhibitor MG-132 (as opposed to the covalent inhibitor Lac) for 3h, after which the inhibitor was removed by changing the medium (washout), and the cells were followed for up to another 2.5h. Both H2a-RFP and CRT-GFP accumulated in the ERQC after treatment, and the ERQC accumulation decreased significantly after washout (Fig. 3A, B). For H2a-RFP, this decrease could be due to recycling to the peripheral ER or alternatively to its proteasomal degradation. However, in the case of CRT-GFP, as it is neither an ERAD substrate nor a short-lived protein, the reduction in its concentration at the ERQC must be ascribed to its recycling back to the peripheral ER.

**Figure 3.**
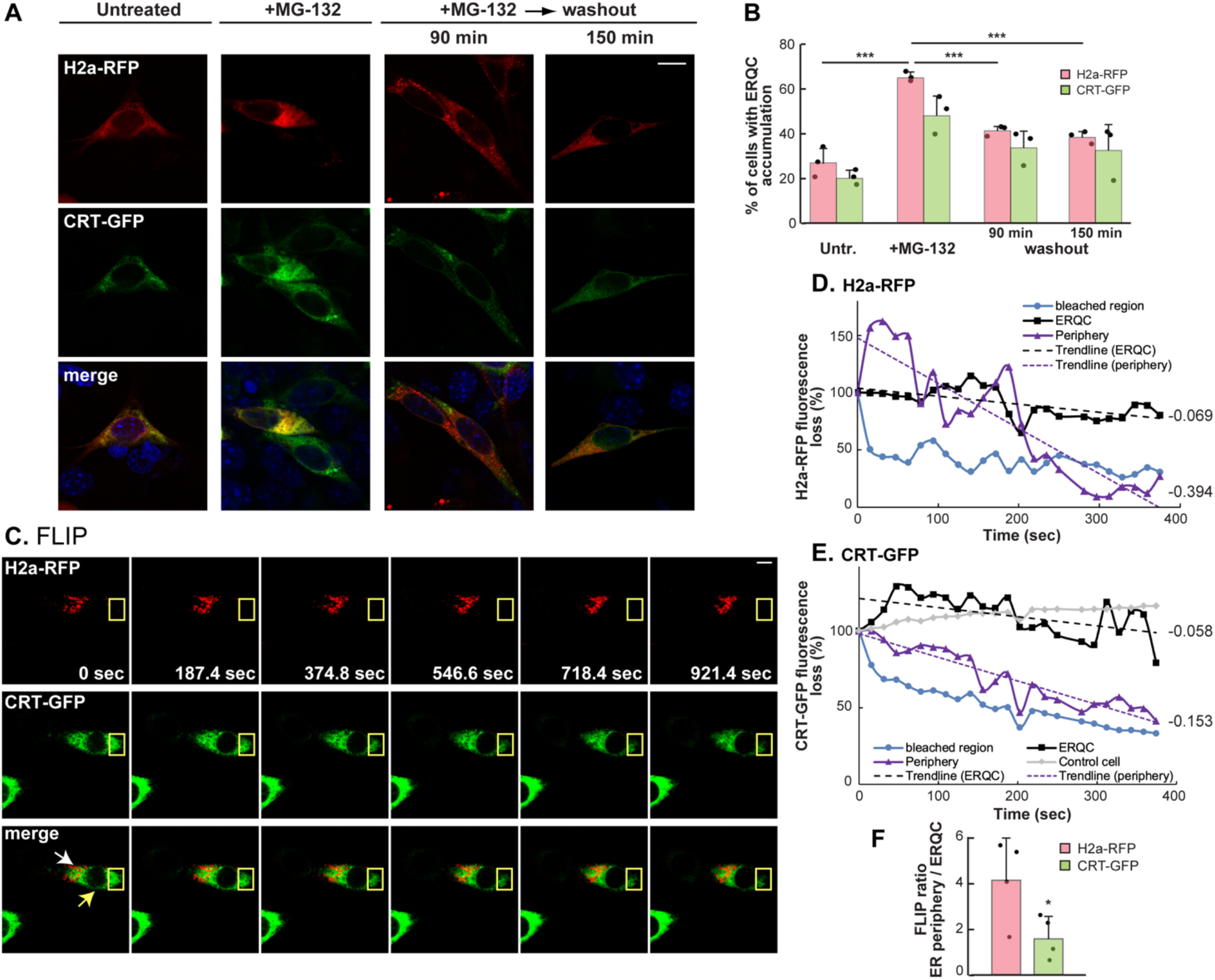
Recycling from the ERQC to the ER is slower than movement within the ER periphery. **(A)** NIH 3T3 cells transiently expressing CRT-GFP and H2a-RFP treated with MG-132 (50μM, 3h), followed by drug washout. The accumulation in the ERQC after treatment reverts to an ER pattern after washout. Bar= 10μm. **(B)** Percent of cells with ERQC accumulation of H2a-RFP or CRT-GFP at the different times, average of 3 independent experiments +-SD. P values for H2a-RFP, +MG-132 vs. untreated = 0.0007, 90 min washout vs. +MG-132 =0.0002, 150 min washout vs. +MG-132 =0.0002. **(C)** Fluorescence loss in photobleaching (FLIP). Lac-treated NIH 3T3 cells (20μM, 3h) were repetitively bleached in an ER periphery region (yellow rectangles) and scanned after each set of bleaching. Fluorescence intensity was measured at the ERQC (white arrow), the ER periphery (yellow arrow) and the bleached region. Representative of 4 experiments. See Suppl. movie S1. **(D-E)** Loss of the fluorescence intensity during FLIP compared to the start point for H2a-RFP (D) and for CRT-GFP (E), in the ERQC, ER periphery, bleached region and in a control cell, during the first 400 sec. Recycling from the ERQC to the periphery is much slower than diffusion in the periphery. Fluorescence in the bleached region was lost quickly, while there was no non-specific bleaching of a nearby but non-irradiated control cell. **(F)** Ratio between the FLIP curve slopes (calculated from linear trendlines) for each protein in the ER periphery compared to the ERQC, average of 4 independent experiments +-SD. P value = 0.049. The relative recycling rate from the ERQC is much slower for H2a-RFP than for CRT-GFP.

To better understand the dynamics of recycling from the ERQC to the peripheral ER we performed a fluorescence loss in photobleaching experiment (FLIP) in live NIH 3T3 cells expressing H2a-RFP and CRT-GFP under continuous proteasomal inhibition. Cells were treated with Lac for 3h for protein accumulation in the ERQC, after which both fluorescent proteins were repetitively bleached in a peripheral region of the ER. H2a-RFP fluorescence intensity in the ERQC declined throughout the time-lapse (Fig. 3C and Suppl. movie S1), though more slowly than in the ER periphery (Fig. 3D). This indicates that H2a-RFP cycles back to the ER periphery, but the nature of the trafficking from the ERQC to the peripheral ER is slower than within the ER, suggesting that it is a vectorial movement, likely vesicular trafficking rather than diffusion. In addition, transient increases in fluorescence intensity in the ER periphery suggest uneven trafficking events, such as that occurring at around 200 sec, with a concomitant decrease in the ERQC region (Fig. 3D). The fluorescence intensity of CRT-GFP in the ERQC also declined more slowly than in the ER periphery (Fig. 3E). Both CRT-GFP and H2a-RFP cycle back from the ERQC to the ER periphery in a non-diffusive manner. To compare the trafficking dynamics of H2a-RFP and CRT-GFP we calculated the ratio between the curve slopes (fluorescence loss rate) for each protein in the ER periphery and in the ERQC (Fig. 3D, E). This ratio was more than 2-fold higher for H2a-RFP than for CRT-GFP when averaging cells from repeat experiments (Fig. 3F), indicating that the movement out of the ERQC compared to its movement within the ER is much slower for H2a-RFP than for CRT-GFP. This could be due to the different fates of the two proteins. Whereas H2a-RFP interacts with ERAD components such as OS-9 at the ERQC ^4,9^, which would cause its retention, no such interactions are known or expected for CRT-GFP, which would be actively shuttling between the ER periphery and the ERQC.

We then looked at another quality control component, ERManI. We had shown that ERManI resides in quality control vesicles (QCVs) at the steady state, concentrating at the ERQC under ER stress ^6^. We performed a FLIP experiment, similar to the one described, in NIH 3T3 cells expressing H2a-RFP and ERManI-YFP under proteasomal inhibition. While H2a-RFP trafficking resembled that of the previous experiment (Fig. 4A, B and Supplementary movie S2), the trafficking of ERManI-YFP was very different than that of H2a-RFP and CRT-GFP. The fluorescence intensity of ERManI-YFP, which appeared in a partially punctate vesicular pattern, decreased rapidly in the ER periphery during the course of the time lapse (Fig. 4C and Fig. 4, Fig. suppl. 1) but barely changed in the ERQC, even after 30 min, double the time of the experiment in Fig. 3C-F, suggesting that it does not recycle back to the ER periphery.

**Figure 4.**
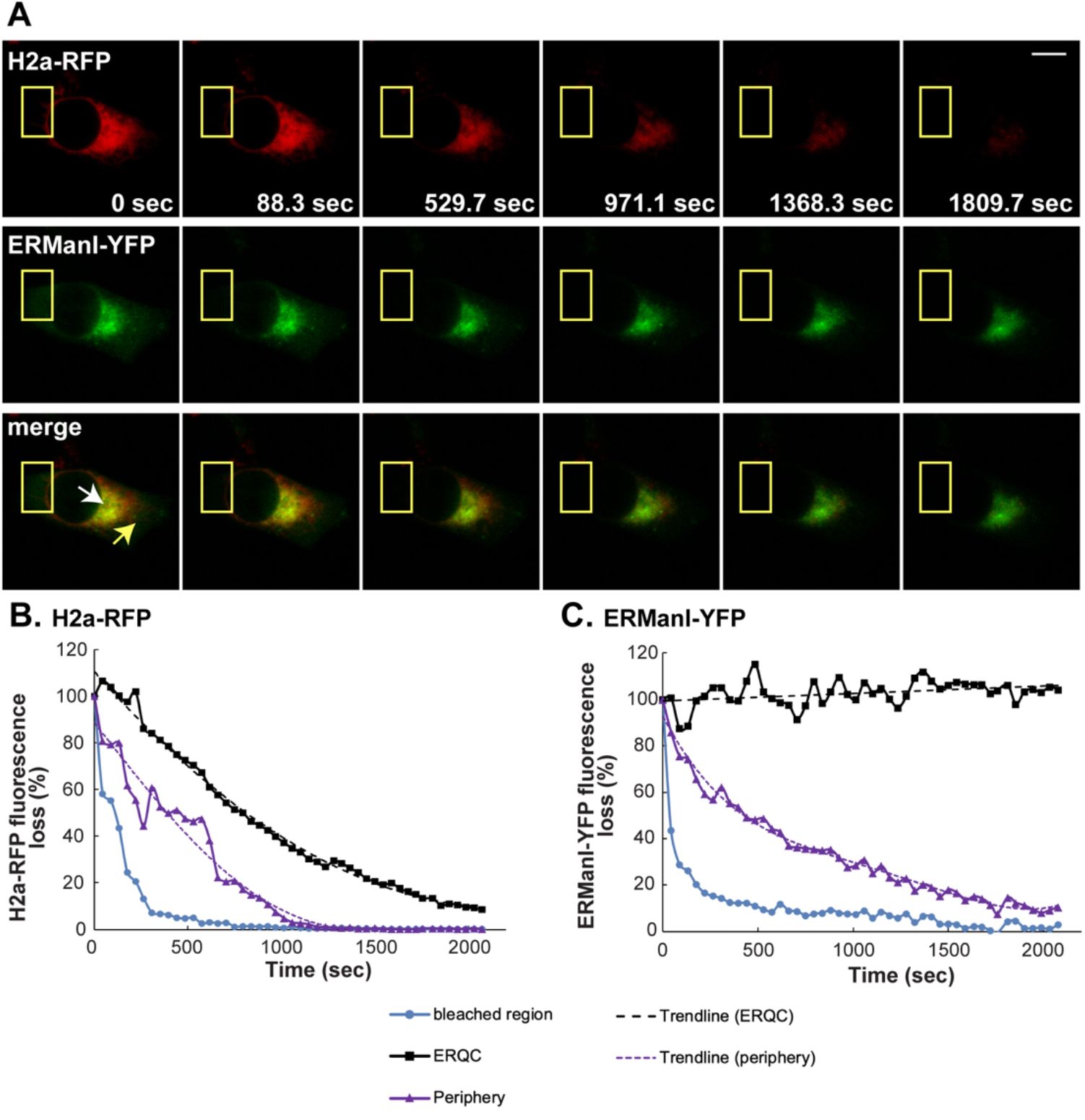
No recycling of ER mannosidase I from the ERQC to the ER. **(A)** Lac-treated NIH 3T3 cells (20μM, 3h) transfected with H2a-RFP and ERManI-YFP expressing plasmids were repetitively bleached in an ER periphery ROI (yellow rectangles) and scanned after each set of bleaching as in Fig. 3C. Bar= 10μm. (**B-C)** The percentage of the fluorescence intensity compared to the start (prior to bleaching) was calculated for each time point for H2a-RFP (B) and for ERmanI-YFP (C), indicating no measurable recycling of ERManI-YFP from the ERQC to the periphery during the time of the experiment. The trendlines in the peripheral region are polynomial (fourth order) and for H2a-RFP in the ERQC it is polynomial of the second order. See Suppl. movie S2.

### Involvement of COPI and COPII coats in the trafficking to the ERQC and back

Given the results of the time lapse experiments, which indicate non-diffusive trafficking from the ERQC to the peripheral ER, consistent with vesicular trafficking, we tested the involvement of vesicular coats. We assessed the effect of overexpression of dominant-negative mutants of ARF1 (ARF1[T31N]) and Sar1 (Sar1[T39N]), which affect COPI and COPII coats respectively ^18,19^, on the localization of H2a-RFP. Similarly to treatment with Bz, ARF1[T31N] caused juxtanuclear accumulation of H2a-RFP (Fig. 5A, B), which was confirmed to be in the ERQC, by comparing with the ERQC localization of OS-9 ^4^ (Fig. 5C, D). In contrast, Sar1[T39N] reduced the number of cells showing spontaneous juxtanuclear accumulation of H2a-RFP (Fig. 5A, B). A coimmunoprecipitation experiment of H2a with OS-9 showed increased interaction in cells overexpressing ARF1[T31N] and conversely, a reduced interaction in the presence of Sar1[T39N] (Fig. 5E, F).

**Fig. 5.**
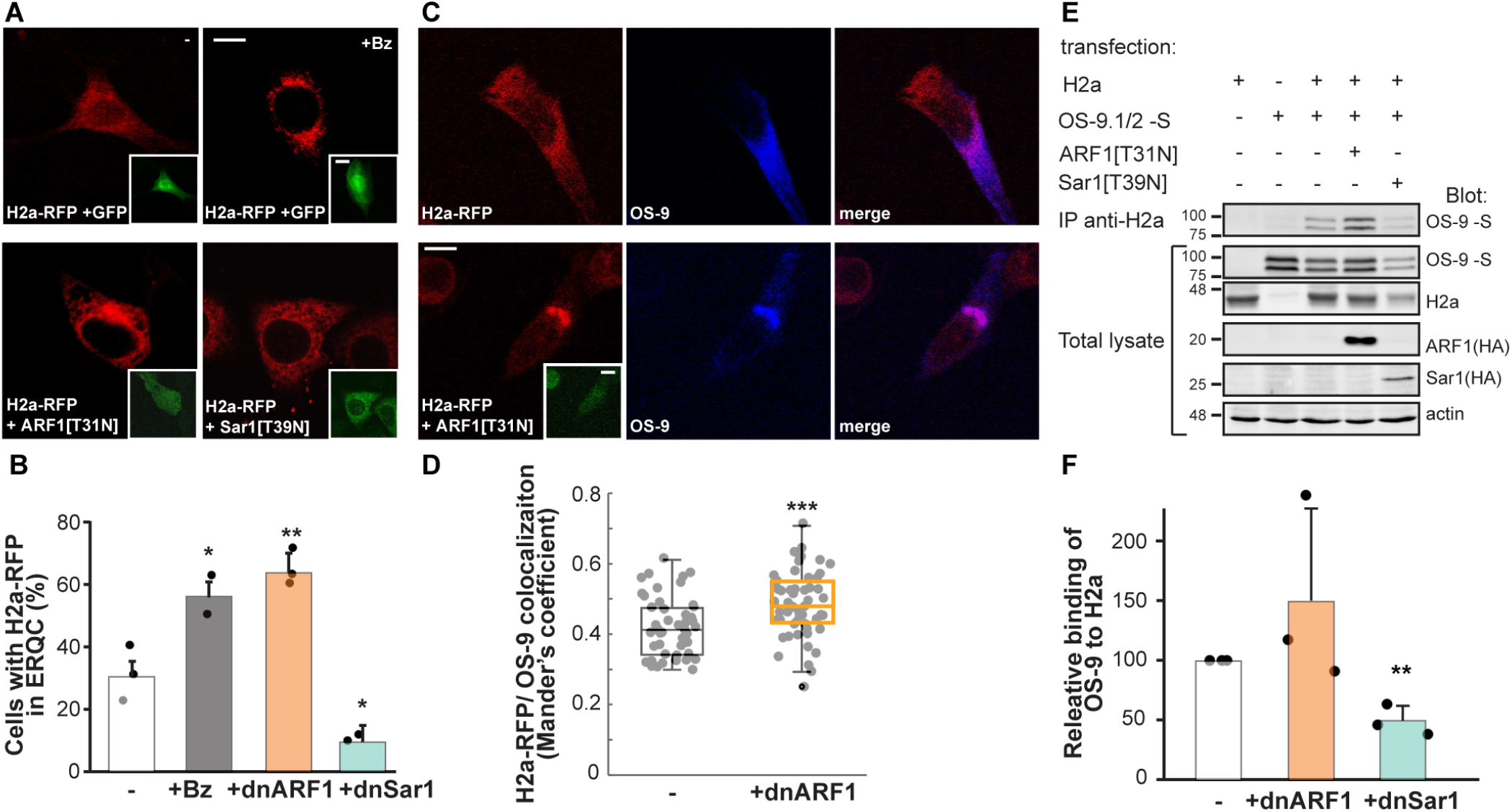
Vesicular coats are involved in trafficking from the peripheral ER to the ERQC and back. **(A-B)** NIH 3T3 cells transiently expressing H2a-RFP together with GFP, ARF1[T31N]-GFP (dnARF1) or Sar1[T39N]-IRES-GFP (dnSar1). Where indicated, cells were treated for 3 h with Bz (1μM). Bar= 10μm. The graph shows percent of cells with ERQC accumulation of H2a-RFP, average of three independent experiments +-SD, which increased +Bz and +dnARF1, and decreased + dnSar1. P values +Bz vs. control = 0.05, +dnARF1 vs. control = 0.005, +dnSar1 vs. control = 0.05. **(C-D)** NIH 3T3 cells expressing H2a-RFP and S-tagged OS-9.1/2, with or without ARF1[T31N]-GFP fixed and stained with anti-S-tag antibody and goat anti-mouse IgG Dy649. Mander’s coefficients indicate that dnARF1 increased H2a-RFP colocalization with OS-9. The graph is a box plot with inclusive medians of 3 independent experiments (~30 cells per experiment); box height=IQR; horizontal bar in the box=median; whiskers extend to the farthest value smaller than 1.5xIQR. P value = 0.0003. **(E)** Immunoprecipitation with anti-H2a of lysates of HEK293 cells expressing H2a together with combinations of OS-9.1/2-S, ARF1[T31N]-HA or Sar1[T39N]-HA, as indicated. Immunoprecipitates and 10 % of the total lysates were immunoblotted with anti-S-tag, anti-H2a, anti-HA and actin (loading control) antibodies. MW markers in kDa are on the left. **(F)** The graph shows increased coimmunoprecipitation of OS-9 with H2a under COPI interference and decreased under COPII interference. Values are relative to control (=100), normalized by total H2a and total OS-9, average of 3 independent experiments +-SD. P value +dnSar1 vs. control = 0.002.

The effect on the compartmentalization by interference with COPII was also studied using iodixanol gradients. Overexpression of Sar1[T39N] caused a shift of H2a from heavier fractions (peaking around fractions 6-7) to lighter fractions (peaking around fractions 4-5) (Fig. 6A, B). There was a reduction in the amount of H2a in the heavier fractions (fractions 6 to 9, which include the ERQC) from 92% of the total protein in the absence of Sar1[T39N] to 41% in its presence (Fig. 6C). To discard the possibility that this is an indirect effect due to the long-term (24h) interference by Sar1[T39N], a similar experiment was done but using short-term (1h) incubation with the PKA inhibitor H89, which inhibits COPII vesicular trafficking ^20^. Cells were incubated with or without Bz (1μM) for 3h with the addition of H89 (50μM) for the last hour. The results were similar to those obtained with Sar1[T39N]. There was a reduction of H2a in ERQC fractions, from close to 100% in untreated cells to 34% in cells treated with H89, and in the presence of Bz from 100% to 60% with H89 (Fig. 6D-G).

**Fig. 6.**
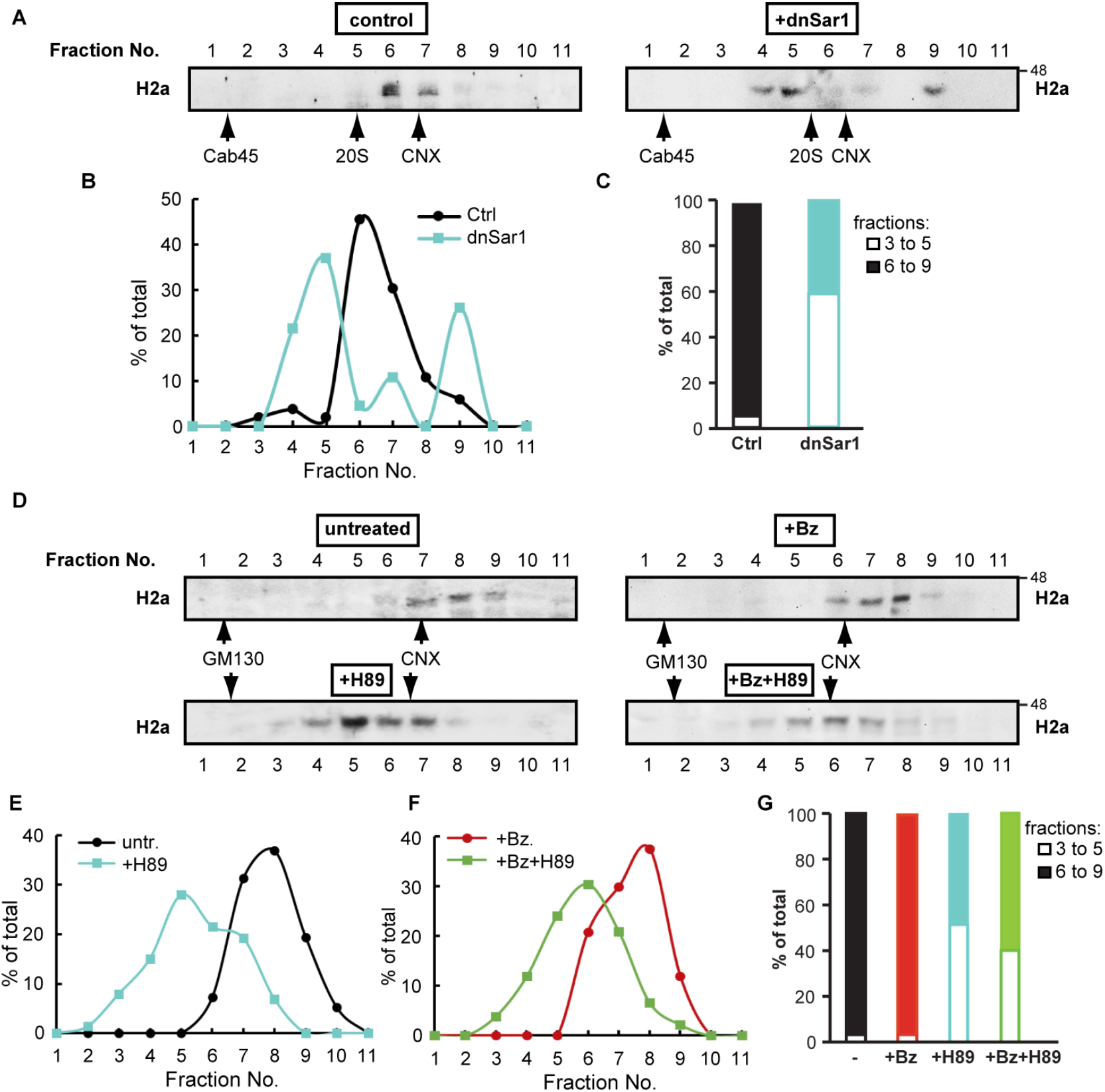
Interference with COPII inhibits ERAD substrate accumulation in the ERQC. **(A)** Homogenates from HEK293 cells expressing H2a with or without Sar1[T39N]-HA (dnSar1) were loaded on an iodixanol gradient (10%-34%) as indicated in Methods. Eleven fractions were collected from the top to the bottom, run on 12% SDS-PAGE and immunoblotted with anti-H2a, anti-Cab45 (Golgi marker), anti-20S and anti-CNX antibodies (peak fractions indicated with arrows). The results shown are representative of 3 independent experiments. **(B)** After quantification, the percentage of total H2a in each fraction was plotted. **(C)** Comparison between the percentage of total H2a in the sum of fractions 3 to 5 and that of fractions 6 to 9 (ERQC) shows that most H2a shifts to lighter fractions under COPII interference by dnSar1. **(D)** Similar to (A), but cells expressing H2a were incubated with 50μM H89 for 1h or 1μM Bz for 3h, or with 1μM Bz for 3h adding 50μM H89 in the last hour of treatment. Anti-GM130 was used here as a Golgi marker. The results shown are representative of 3 independent experiments. (**E-F)** The intensity of each band was quantified and the percentage of total H2a in each fraction was plotted and compared between the control and +H89 (E), or +Bz and +Bz+H89 (F). **(G)** Comparison between the percentage of total H2a in the sum of fractions 3 to 5 and that of fractions 6 to 9 (ERQC) shows that H2a shifts to lighter fractions under COPII interference by H89 treatment.

It is important to note that in no case was there any detectable ERAD substrate in Golgi fractions (peaking around fractions 1-2 as indicated by the Golgi markers Cab45 or GM130, Fig. 6 and Fig. 6, Fig. suppl. 1).

Altogether these results suggest that COPII is necessary for trafficking of the ERAD substrate from the peripheral ER to the ERQC and interaction with OS-9, whereas COPI appears to facilitate recycling from the ERQC to the peripheral ER.

### COPI and COPII are involved in the regulation of misfolded protein targeting to retrotranslocation and ERAD

The requirement of COPII function for efficient interaction of the ERAD substrate with the HRD1-SEL1L-associated OS-9 suggested that it may also regulate its targeting to retrotranslocation and proteasomal degradation. We tested the effect on retrotranslocation using the BAP/BirA system, developed by the group of Burrone ^21^. This system utilizes the E. coli biotin ligase BirA expressed in the cytosol of mammalian cells to specifically biotinylate a 15 amino acid biotin-acceptor peptide (BAP). Secretory proteins with a BAP tag on their ER luminal side will be biotinylated only after retrotranslocation. We made a construct to express H2a with BAP attached to its luminal C-terminus (H2a is a type II membrane protein). H2a-BAP was biotinylated poorly when coexpressed with cytosolic BirA and to a much higher extent when coexpressed with a luminally expressed secBirA, which contains a signal peptide for translocation into the ER ^21^ (Fig. 7A). Biotinylated H2a-BAP in the presence of cytosolic BirA represents retrotranslocated H2a-BAP molecules. Interestingly, these biotinylated molecules concentrated at the ERQC fractions in iodixanol gradients (especially upon proteasome inhibition), with only a minor fraction at light densities, suggesting that H2a molecules remain associated to the cytosolic side of the ERQC membrane prior to degradation (Fig. 7B).

**Fig. 7.**
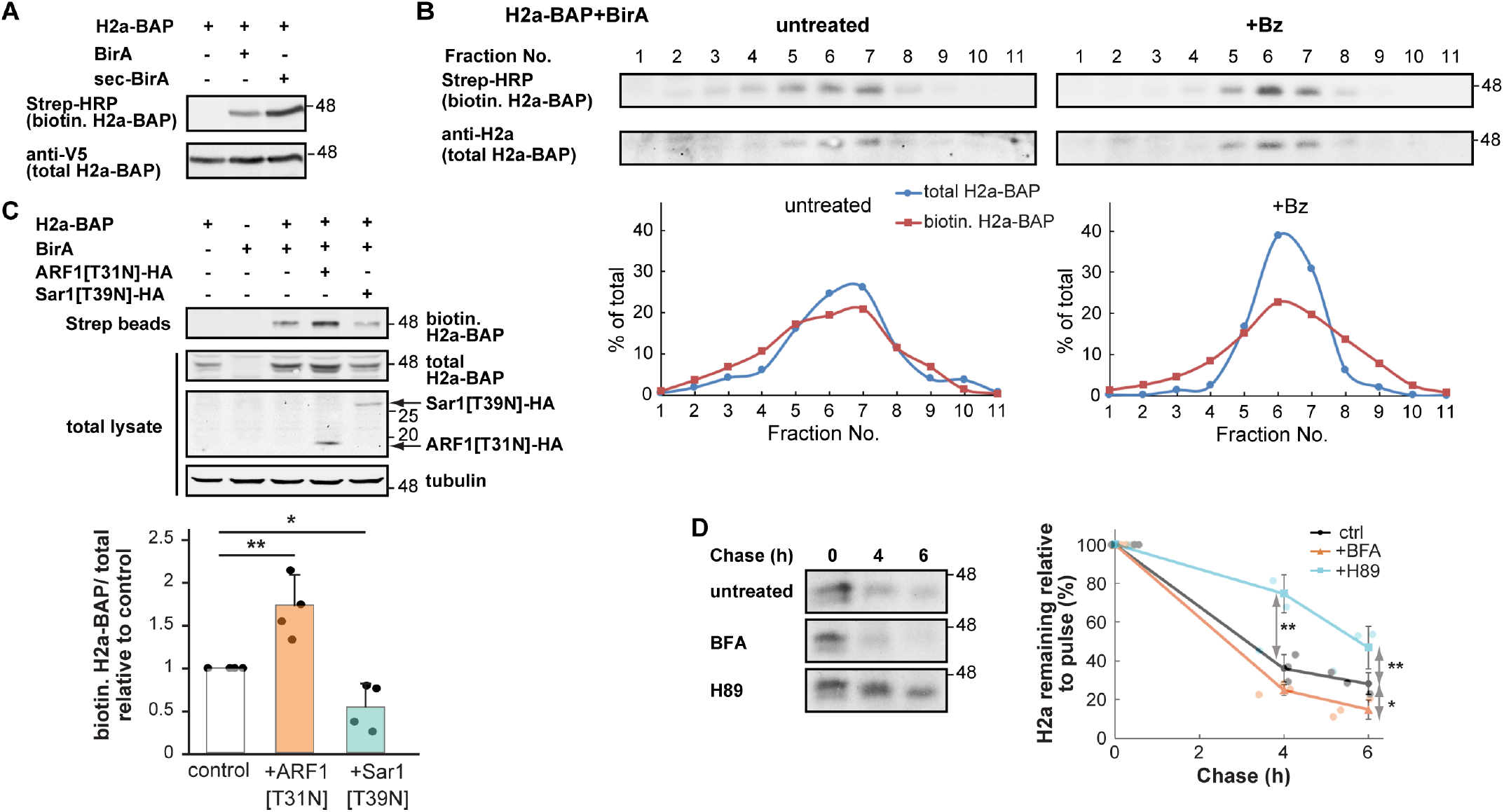
COPII-dependent movement from the peripheral ER to the ERQC is required for ERAD substrate retrotranslocation and degradation. **(A)** HEK293 cells transiently expressing H2a-BAP with or without a cytosolic biotin ligase, BirA, or an ER luminal one, Sec-BirA. Cells were incubated 24h post-transfection with biotin (100μM) for 30 min before lysis. Cytosolic BirA biotinylates BAP exposed on the cytosolic side of the ER upon retrotranslocation of H2a-BAP, while Sec-BirA biotinylates luminal H2a-BAP. Strep-HRP was used to detect biotinylated H2a-BAP and anti-V5 to detect total H2a-BAP. MW markers (kDa) on the right. **(B)** Fractions of iodixanol gradients from cells expressing H2a-BAP and BirA, were immunoblotted as in (A), (with anti-H2a instead of anti-V5). Both retrotranslocated and total H2a-BAP peak in fractions 6-7, which include the ERQC. **(C)** Biotinylated proteins from cells expressing H2a-BAP together with BirA, ARF1[T31N]-HA or Sar1[T39N]-HA, (biotin for 3h), were precipitated with streptavidin-agarose. Precipitates and 10% of total cell lysates were immunoblotted with anti-H2a, anti-HA-tag or anti-tubulin antibodies. Retrotranslocated (biotinylated) H2a-BAP increased almost two-fold under COPI interference and was reduced to half under COPII interference. Values are averages of 4 independent experiments ±SD, relative to total H2a-BAP and standardized by the control (=1). P value +dnARF1 vs. control = 0.007, +dnSar1 vs. control = 0.02. **(D)** Pulse-labeling for 20 min with [^35^S]cysteine and chase for the indicated times with or without 5μg/ml BFA or 100μM H89. H2a degradation (percentage remaining relative to pulse) increased under COPI interference (+BFA) and was strongly inhibited under COPII interference (+H89). The graph shows an average of 3 independent experiments ±SD. P value 6h +BFA vs. control = 0.03, 6h +H89 vs. control = 0.01, 4h +H89 vs. control = 0.003.

We then tested the effect of ARF1[T31N]-HA and Sar1[T39N]-HA on retrotranslocation. The fraction of retrotranslocated H2a-BAP increased approximately 2-fold under COPI interference (+ARF1[T31N]-HA) and decreased to about half under COPII interference (+Sar1[T39N]-HA) (Fig. 7C). This suggests that COPII is required for efficient targeting of the ERAD substrate to retrotranslocation. Concerning COPI, its role in recycling from the ERQC to the peripheral ER, which we have observed in the previous experiments, could reduce the availability of molecules for retrotranslocation. However, if COPI would be involved in events taking place after retrotranslocation, this could also explain the increase seen by interfering with COPI.

As retrotranslocation is a rate-limiting step in ERAD, we reasoned that interference with COPI and COPII may affect the degradation rate of the ERAD substrate. We tested this by pulse-chase analysis, to examine H2a degradation over time under COPI/COPII interference. HEK293 cells transiently expressing H2a were metabolically labeled with [^35^S]cysteine. To minimize indirect effects of long-term expression of the dominant negative constructs, we decided in this case to incubate cells only during the chase period of up to 6h with brefeldin A (BFA) (5 μg/ml), a known inhibitor of COPI vesicular transport, or H89 (100 μM). In untreated cells, H2a degraded with time, and 28% of the labeled molecules remained after 6h of chase (Fig. 7D). COPII interference with H89 inhibited the degradation, with 47% of the labeled protein remaining after 6h chase. Interestingly, COPI interference with BFA accelerated the degradation, with 15% remaining after 6h, a reduction to almost half of the untreated control.

Altogether, these results suggest involvement of COPII in the targeting of the ERAD substrate to retrotranslocation and degradation. In the case of COPI, the increased degradation of the ERAD substrate seen by interfering with its function suggests that it does not participate in events taking place after retrotranslocation, but rather it functions in the recycling from the ERQC to the peripheral ER, which reduces the availability of ERAD substrate molecules both for retrotranslocation and ι ilfimatαl\/ fnr thαir rlαnrarlatinn /’Pin R\

## Discussion

Previous work in our lab had characterized the ERQC as a distinct compartment, localized in the centrosomal region of the cell and not colocalizing with the Golgi apparatus, ERGIC or other organelles ^2,3^. OS-9 resides constitutively in this compartment ^4,22^, but ERAD substrates and other machinery components are observed to concentrate at the ERQC only upon accumulation of ERAD substrates, by inhibiting their proteasomal degradation (as done here), by their overexpression or by induction of Herp, downstream of the activation of the PERK pathway ^3,4^. For example, recruitment to the ERQC can be achieved by overexpressing a phosphomimetic mutant of eIF2α(S51 D) ^3^, inhibiting eIF2α-P dephosphorylation or overexpressing Herp ^4^. We had concluded from those results that even in unstressed conditions, a transient accumulation of unfolded protein molecules causes a temporary activation of the PERK pathway and an induction of Herp, leading to dynamic protein recruitment to the ERQC. Therefore, ERAD machinery components are recruited according to the demand for protein disposal and ERAD complexes might be transiently assembled only when needed, followed by disassembly and dispersal in an “ERAD cycle”. We have now observed that also inhibition of COPI vesicular trafficking with a dominant-negative mutant of ARF1 causes an accumulation of the ERAD substrate in the ERQC (Fig. 5), suggesting that the recycling or dispersal is inhibited. Many quality control and ERAD components are recruited to the ERQC, also from the cytosolic side, including proteasomes, but some are excluded, such as the abundant chaperone BiP (Fig. 1)^5^.

Both glycoproteins and non-glycosylated ERAD substrates concentrate at the ERQC, as they share much of a common quality control machinery ^1,7^. However, we have observed that the trafficking of CRT to the ERQC is dependent on the presence of an active lectin domain (Fig. 2), and abrogated in a mutant without lectin activity ^17^, suggesting that it follows the accumulation of its glycoprotein clients. CRT appears to cycle actively between the peripheral ER and the ERQC (Fig. 3). In contrast, another glycoprotein quality control component, ERManI, does not recycle (Fig. 4) and seems to be effectively sequestered in QCVs or in the ERQC.

We have described the QCVs as a new type of vesicles, where the mannosidases ERManI and mannosidase IA (ManIA) are normally kept segregated from their substrate glycoproteins ^6,23^. Only occasionally do the glycoproteins interact with the mannosidases at the QCVs or at their fusion sites with the ER. The QCVs are dependent on COPII, but coat components were not observed on the vesicles. Future studies should determine whether the QCVs are the same carriers for the trafficking of the ERAD substrates.

Interference with COPII inhibits ERQC localization of the ERAD substrate and reduces its interaction with OS-9, retrotranslocation and degradation (Fig. 5-8). One possibility is that the interference with COPII affects the QCVs, thus causing an inhibition of substrate interaction with the mannosidases, which reduces mannose trimming and delivery to ERAD. However, the slow dynamics of H2a-RFP to the ERQC (Fig. 2) and its slow recycling (Fig. 3-4) suggest vesicular transport of the substrate and not an indirect effect of the mannose trimming. Trafficking of ERAD substrates and recruitment of machinery components at the ERQC is also microtubule dependent ^2^.

**Fig. 8.**
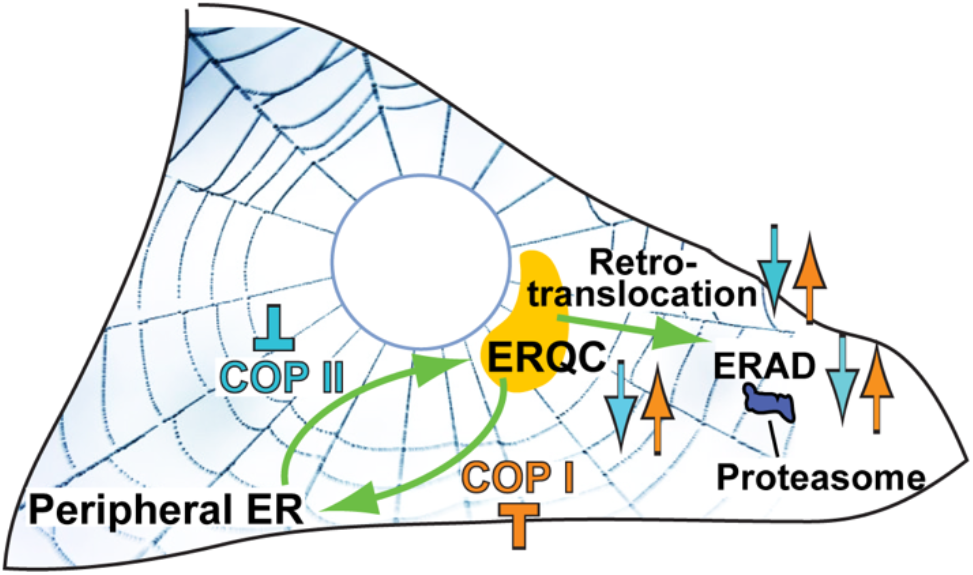
Model of COPI and COPII involvement. COPII is involved in the movement of ERAD substrates from the peripheral ER to the ERQC. Under COPII interference this movement is inhibited. The ERAD substrates are then mostly scattered in the peripheral ER instead of accumulating in the ERQC. The retrotranslocation is then decreased, followed by a slower degradation rate (light blue arrows). The recycling of ERAD substrates from the ERQC to the ER is dependent on COPI. COPI interference hinders this recycling, causing accumulation of ERAD substrates in the ERQC. The retrotranslocation is then increased and the degradation of the substrates is accelerated (orange arrows).

The COPII-dependent sorting to an ER subdomain, named ER-associated compartment (ERAC), was found to be required for proteasomal degradation of an ERAD substrate, mutant CFTR, in *S. cerevisiae.* Perhaps the ERAC is a functional homolog of the ERQC in yeast, although there are some differences, such as the accumulation of the BiP homolog, Kar2 in the ERAC whereas BiP does not accumulate in the ERQC ^24,25^.

COPI and II vesicular trafficking is traditionally associated to movement between the ER and the Golgi. However, COPII coats have recently been implicated in other pathways as well. For example, in the formation of ER whorls, which appear in certain conditions and were found to be PERK and COPII dependent ^13^. COPII was also found to be involved in the initiation of the autophagic process in yeast ^14,15^ and mammalian cells ^26^. Some interesting recent reports depart from the classical model, suggesting that COPII concentrates at the ER exit sites, creating a boundary, but does not coat the vesicular carriers ^27,28^. COPII and also COPI involvement might be in the vesicle budding stage, with alternative factors possibly targeting the vesicles to different locations ^29,30^.

There are reports of misfolded proteins that are sent back to the ER from the Golgi ^31^ or from the ERGIC ^32^. However, in the case of H2a that we have analyzed here, it is retained in the ER and does not traffic to the Golgi nor to the ERGIC ^4,5,7^.

Our results show that by interfering with misfolded protein delivery to the ERQC (Fig. 5, 6), interference with COPII surprisingly inhibits retrotranslocation and ERAD (Fig. 7, 8). COPII mutations are reported to cause ER stress, which has been interpreted as resulting from the accumulation of secretory proteins that cannot travel to the Golgi ^33^, but an additional cause might be an outcome of ERAD inhibition.

## Materials and Methods

### Materials

Streptavidin-agarose-beads, ALLN, MG132, Bz, H89 and biotin were from Sigma. Lac was from Calbiochem. Protein A-Sepharose was from Repligen (Needham, MA). Other common reagents were from Sigma-Aldrich.

### Antibodies

Rabbit polyclonal anti-H2a amino-terminal and carboxy-terminal antibodies were the one used in previous studies ^34^. Rabbit polyclonal anti-Cab45 was described in ^35^ and anti-BiP was the one used before ^4^. Rabbit polyclonal anti-GM130 was from BioLegend, anti-CNX from Sigma and anti-dsRED from MBL. Rabbit polyclonal anti-19S proteasome from Thermo scientific and anti-20S proteasome from Abcam and were kind gifts from Michal Sharon, Weizmann Institute.

Mouse monoclonal anti-HA was from Sigma or from BioLegend, anti-actin and anti-ß-tubulin from Sigma, anti-S-tag from Novagen, anti-V5 from GenScript and anti-GFP from Santa Cruz Biotechnology. Strep-HRP, goat anti-mouse IgG-HRP, goat anti-rabbit IgG-HRP and goat anti-mouse IgG Dy649 were from Jackson-Immuno-Research Labs.

### Plasmids and constructs

psmd14-YFP was a kind gift of A. Stanhill, (Open Univ., Israel)) ^36^. CRT-GFP and mutCRT-GFP: CRT and CRT(Y108F) expressed in pEGFP-C1 were kind gifts from Erik Snapp, (Janelia) ^17^. H2a in pcDNA1 was described before ^2^. BiP-GFP, H2a-RFP, S-tagged-OS-9.1 and S-tagged-OS-9.2 in pcDNA3.1(+) were used before ^3,4,6,23^. ERManI-YFP was constructed with human ERManI cloned into peYFP-N1 (Clontech) using Hind III and Xma I yielding ERManI fused on its C-terminus to YFP through a 6 amino acid flexible linker (SGGGGS). Sar1[T39N]-HA in pTARGET or in IRES-GFP (Clontech) were used before ^6^. ARF1[T31N]-HA in pcDNA3.1(+)^37^, ARF1[T31N]-GFP ^38^ and pBABE-puro were from Addgene. BirA and Sec-BirA plasmids were a kind gift from Gianlucca Petris and Oscar Burrone (ICGEB, Trieste, Italy) ^21^. H2a-BAP was cloned in pcDNA3.1 using restriction enzymes EcoR1 and XbaI for H2a(G78R) followed by a C-terminal BAP-SV5 tag using XbaI ^39^.

### Cell culture, media and transfections

NIH 3T3, CHO and HEK293 cells were grown in DMEM supplemented with 10% bovine calf serum at 37 °C under 5% CO2. Transfections of NIH 3T3 and CHO cells were carried out using a Neon MP-100 microporator system (Life Technologies, Carlsbad, CA). Transfection of HEK293 cells was performed using the calcium phosphate method. Stable CHO cell lines expressing CRT-GFP or CRT(Y108F)-GFP were obtained by co-transfection with pBABE-puro (Addgene) and selection with puromycin.

### Microsome isolation and gradient fractionation

Cells were homogenized by passing through a 25-gauge needle 5 times followed by 30 strokes with a Dounce homogenizer (Kontes glass co.) in an iso-osmotic buffer consisting of 10 mM Hepes (pH 7.4) and 250 mM sucrose. Debris and nuclei were pelleted by centrifugation at 1,000xg at 4°C for 10 minutes. Then the supernatants were loaded on top of a iodixanol gradient (10 to 34%) as previously described ^23^. The gradients were ultra-centrifuged at 24,000 rpm (~98,500g, Beckman SW41 rotor) at 4°C for 16 hours. Eleven fractions were collected from top to bottom and subjected to gel electrophoresis followed by immunoblotting.

### Metabolic labeling

Subconfluent (90%) cell monolayers in 60-mm dishes were labeled for 30 min with [^35^S]Cys and chased for different periods of time with normal DMEM plus 10% FCS, lysed, and immunoprecipitated with anti-H2a carboxy-terminal antibody as described previously ^23,34^. Quantitation was performed in a Fujifilm FLA 5100 phosphorimager (Tokyo, Japan).

### Immunoprecipitation and Immunoblotting

Cell lysis and immunoprecipitation methods are described in ^9^. For precipitation with streptavidin-agarose beads, cell lysates were incubated with the beads on a vertical rotating mixer at 4°C overnight. After centrifugation, the beads were washed three times with 0.5% Triton X-100, 0.25% NaDOC and 0.5% SDS in PBS and once with PBS. Samples were boiled in sample buffer for 5 minutes and run on SDS-PAGE under reducing conditions. Transfer to a nitrocellulose membrane, Immunoblotting, detection by ECL, and quantitation in a Bio-Rad ChemiDocXRS Imaging System (Hercules, CA) were done as described previously ^22^. Quantity one or ImageJ software were used for the quantitation.

### BAP construct probing

BAP-tagged H2a (H2a-BAP) was transfected and probed as described before ^39^. Briefly, H2a-BAP was cotransfected in HEK293 cells with an equal amount of plasmids carrying either BirA (cytosolic biotin ligase) or Sec-BirA (ER luminal biotin ligase). Biotin (100μM) was added 24 hours post transfection for 30 to 180 min as indicated, cells were lysed, subjected to SDS-PAGE and blots reacted with Strep-HRP in PBS-TWEEN (0.5%), followed by ECL detection. Total H2a-BAP was detected with anti-H2a amino-terminal or anti-V5 antibodies.

### Immunofluorescence Microscopy

Immunofluorescence was performed as described previously ^2,16,40^. Briefly, cells grown for 24 h after transfection on coverslips in 24-well plates were fixed with 3% paraformaldehyde for 30 min, incubated with 50mM glycine in PBS, and permeabilized with 0.5% Triton X-100. After blocking with normal goat IgG in PBS/2% BSA, they were exposed to primary antibody for 60 min, washed and incubated for 30 min with a secondary antibody, followed by washes and staining of nuclei with DAPI. Specimens were observed with a Leica DMRBE microscope, or confocal microscopy on a Zeiss laser scanning confocal microscope (LSM 510; Carl Zeiss, Jena, Germany). ImageJ was used to quantify fluorescence intensity and to calculate Pearson’s and Mander’s coefficients (using JACOP) for colocalization studies.

### Live cell imaging

Cells were transfected and grown in 35mm culture dishes on 25mm coverslips. 24 hours post-transfection images were captured using a Zeiss LSM 510 Meta confocal microscope. Cells were kept in a stage incubator which provides tissue culture conditions (37°C, CO2). For FLIP experiments, GFP bleaching was achieved with an argon laser at a wavelength of 488nm, 100% output. RFP bleaching was achieved with solid state laser at a wavelength of 561nm, 15% output. Bleaching was done repetitively in an ER periphery region and scanned after each set of bleaching (200 iterations at each time point). Analyses were performed in ImageJ software.

### Statistical Analysis

The results are expressed as average ± SD or mean ± SEM as indicated. Student’s t-test (two-tailed) was used to compare the averages of two groups. Statistical significance was determined at P<0.05 (*), P<0.01 (**), P<0.001 (***).

## Acknowledgements

We would like to thank Erik Snapp, Ariel Stanhill, Michal Sharon, Gianlucca Petris and Oscar Burrone for plasmids and reagents. Work was supported by grants (1593/16 and 2577/20) from the Israel Science Foundation (GZL).

