## Supplementary material for "COP I and II dependent trafficking controls ER-associated degradation in mammalian cells": Suppl. figures

**Supplementary figures**

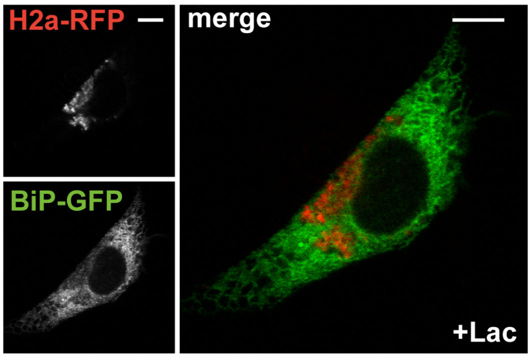


**Fig. 1, Fig. suppl. 1. BiP-GFP does not accumulate at the ERQC upon proteasomal inhibition.** Fluorescence microscopy of NIH 3T3 cells treated with Lactacystin (Lac, 20μM, 3h), transiently expressing H2a-RFP and BiP-GFP. H2a-RFP concentrates at the juxtanuclear ERQC, whereas BiP-GFP does not. There is almost no colocalization of H2a-RFP and BiP-GFP. Bars= 10μm.

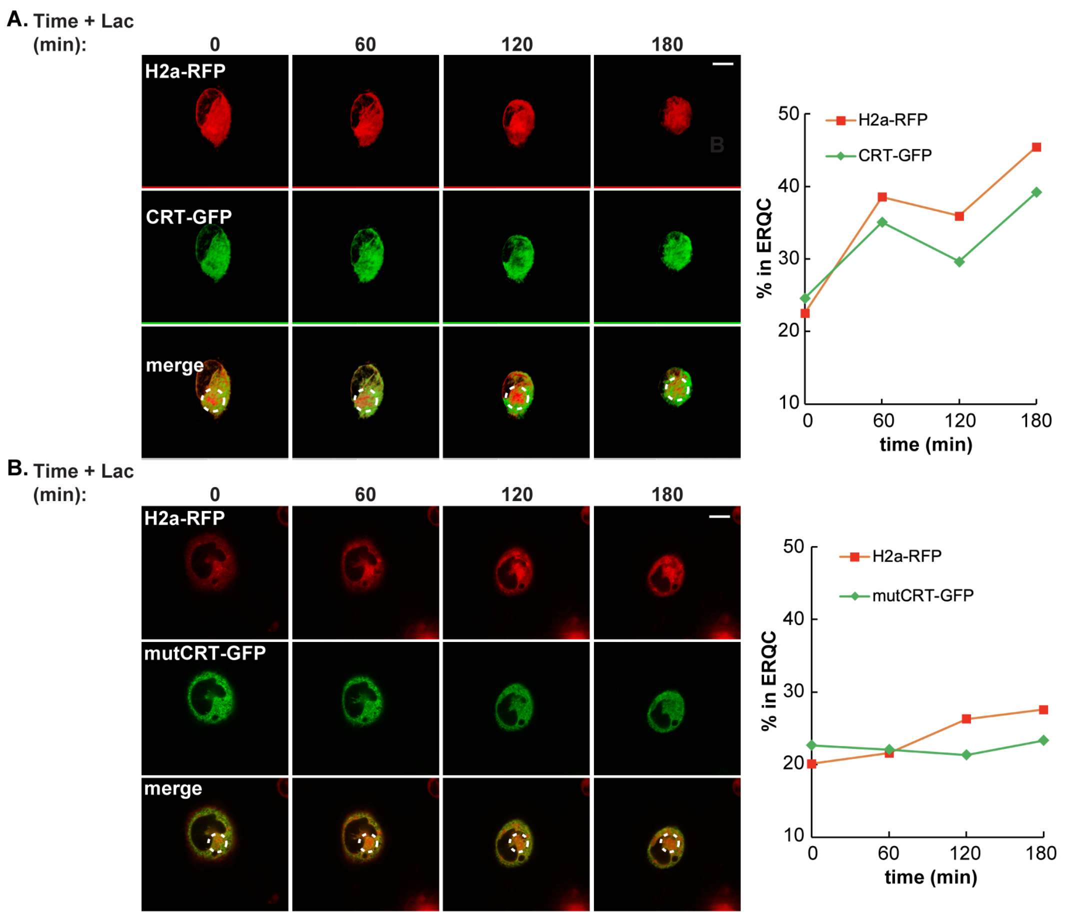


**Fig. 2, Fig. suppl. 1. Movement of H2a-RFP and CRT-GFP to the ERQC and dependence of CRT-GFP on lectin activity.** Similar to Fig. 2 (A-D), a repeat experiment is shown. Bars= 10μm.

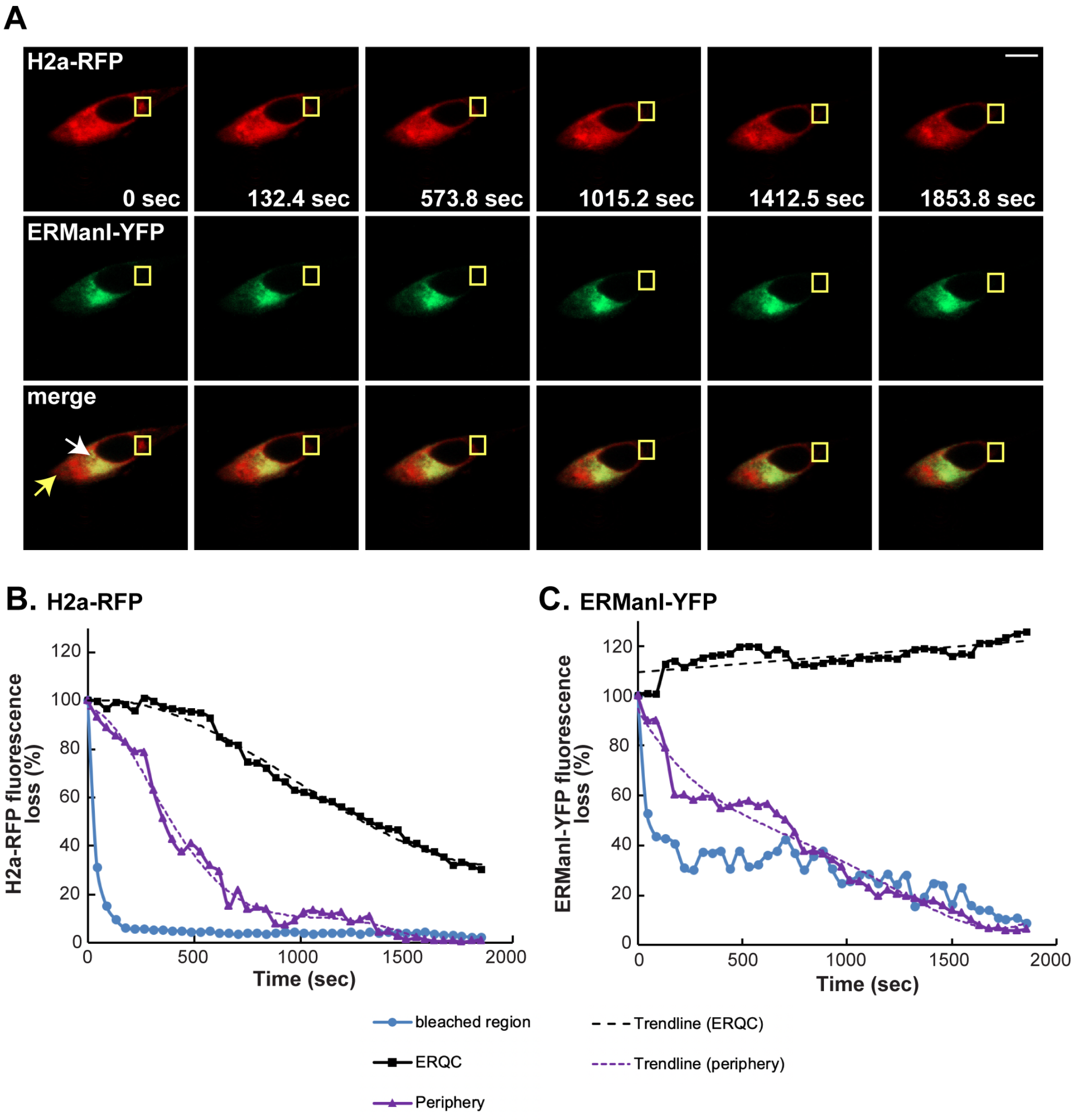


**Fig. 4, Fig. suppl. 1. No recycling of ER mannosidase I from the ERQC to the ER.** Similar to Fig. 4, a repeat experiment is shown.

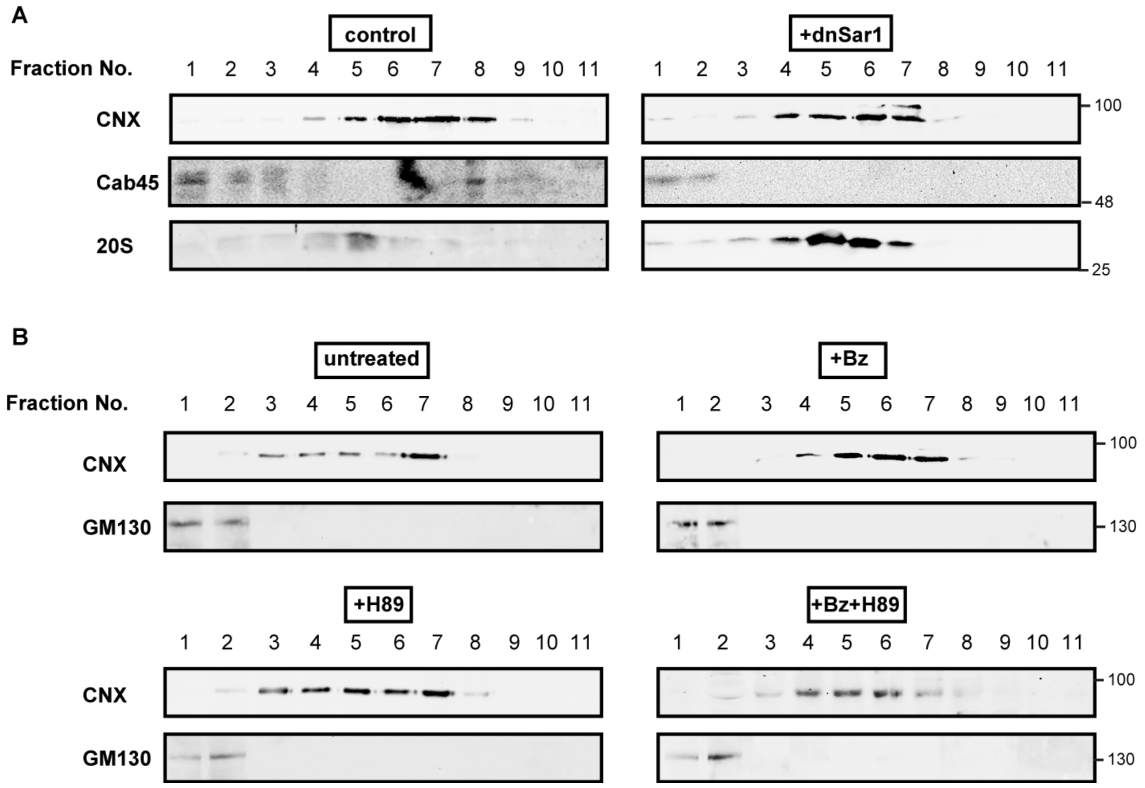


**Fig. 6, Fig. suppl. 1. Organelle markers under COPII inhibition.** Iodixanol gradients of the markers indicated with arrows in Fig. 6. Whereas the markers of the Golgi and 20S proteasomes did not show any significant change in migration, CNX (which localizes partially to the ERQC) shifted slightly to lighter fractions upon COPII inhibition by dnSar1 (A) or H89 treatment (B).

**Supplementary movies**

**Movie S1.**
FLIP of CRT-GFP and H2a-RFP as described in Fig. 3 C-F.

**Movie S2.**
FLIP of ERManI-YFP and H2a-RFP as described in Fig. 4.
